# A low-cost, AI-powered Measurement Verification and Reporting System for growing trees with smallholder farmers

**DOI:** 10.1101/2023.11.29.569237

**Authors:** Edward Idun Amoah, Peter McCloskey, Rimnoma Serge Ouedrago, John Chelal, Chelsea Akuleut, Binti Ibrahan Mwambumba, Brian Kipchirchir Meli, Christabel Akinyi Oyudi, Edna Santa Kibwamga, Eunice Kwamboka Cleophas, Fredrick Odhiambo Ochola, Catherine Njeri Wangiru, Kelvin Morang’a, Lyon Wilson Mushira, Maureen Kalegi Maboke, Nancy Syonthi Titus, Serah Lanoi Oltimbao, Sheilah Awour Odawa, David Peter Hughes

## Abstract

Limited access to low-cost tools to measure, report, and verify (MRV) tree growth with smallholder farmers limits the scaling of tree planting efforts in developing countries. Artificial Intelligence (AI) offers the potential for low-cost, reliable, and accessible measurement and verification tools to be developed for an MRV platform to scale tree planting efforts in developing countries. Here, we present an AI-powered non-contact tree diameter measurement and verification tool. We have developed an AI-powered algorithm that accurately estimates the diameter of a tree from an image of the tree with a reference object. This non-contact measurement method utilizes semantic segmentation and image processing techniques to analyze an image of the tree with the reference object. The performance of the proposed method was evaluated on 142 trees with tape-measured diameters at breast height ranging from 5 to 60 cm. A regression analysis between predicted and measured diameter values had an R^2^ and an RMSE of 0.97 and 2.23 cm, respectively. Thus, using a smartphone application, the non-contact method developed here can empower anyone to accurately measure and report tree growth by just taking pictures of the trees with the reference object. The images submitted with on-farm measurements serve as data for future verification operations using the AI-powered algorithm. With the reference object serving as a unique tree identifier, a tree’s survival and diameter measurements can be tracked over time. The MRV system described here, with the developed AI-powered non-contact tree diameter measurement and verification tool, can empower organizations to plant, grow, and monitor trees with anyone, including smallholder farmers.

## 1 Introduction

Tree planting remains the most effective strategy to mitigate climate change (Bastin et al., 2019). While tree planting efforts have received support from many organizations and institutions, these programs rarely monitor tree growth and survival once the trees have been planted. In programs where the trees have been monitored, it was discovered that tree survival was extremely low. One study reported a survival rate of less than 10% on more than half of the sites where trees were planted for a tree planting program in Sri Lanka (Kodikara et al., 2017). A planted tree can only capture CO^2^ if the tree is well-maintained and survives over time. Tree planting efforts for climate mitigation require long-term targets, including maintenance and monitoring of planted trees over time. This is especially important under climate change, which poses severe challenges for some species but could also accelerate the growth of other species (Fisichelli et al., 2014; Menezes-Silva et al., 2019).

Tree diameter monitoring is one of the primary methodologies for monitoring tree biomass growth in Ecology. The global network of 59 long-term forest dynamic research sites (CTFS-ForestGEO) in 24 countries, monitored as early as 1981, has been critical to helping study forest responses to global change (Anderson-Teixeira et al., 2015). In these sites, a standard protocol was established where the diameters at the breast height (dbh) of all identified tree species with a diameter ≥ 1 cm were measured at regular temporal intervals of 5 years, allowing a precise characterization of the forest structure, diversity, and dynamics over space and time (Anderson-Teixeira et al., 2015). The measurement and monitoring of trees require skilled personnel and can be costly (Weiskittel et al., 2015). This challenge can be a bottleneck for tree monitoring efforts, especially in developing countries with limited resources.

Non-contact measurement methods can potentially address the major fallbacks of contact measurement methods while delivering comparable performance in accuracy. Diameter measurement methods can be divided into two types: (1) contacting methods with tools like calipers and measuring tapes and (2) non-contact methods with a series of innovative technologies (Clark et al., 2001; Shao et al., 2022). The non-contact measurement methods can be further divided into (1) reference-based and (2) non-reference methods. The reference-based measurement method procedure involves identifying the boundaries of a tree and reference object in an image and then using the reference object to tree ratio for diameter estimation. One major drawback of this method is the lack of reliable identification of tree trunks and reference objects under various conditions like lighting variations (Megalingam et al., 2022; Shao et al., 2022). Various methods for non-reference methods have been developed, but these methods can be expensive to implement. Extra sensors like LiDAR and complex data processing software like point cloud analysis are required for non-reference measurement methods (Koreň et al., 2017; Woo et al., 2021; Tran et al., 2023; Shao et al., 2022). Non-reference measurement methods can provide a robust method of measurement. However, their cost of implementation renders these methods inaccessible for smallholder farmers.

In this study, we have developed a non-contact reference-based method for tree diameter measurements and verification. A state-of-the-art deep-learning semantic segmentation model was fine-tuned for reliable and accurate tree and reference object identification. This method can accurately estimate tree diameter by leveraging just an image of the tree with a tree tag reference object. The AI-powered algorithm developed in this study can serve as a measurement and verification tool for an MRV platform to monitor tree survival and growth with anyone with access to a smartphone. The reference object used in this system serves as a unique tree identifier, which enables tree survival and growth to be monitored over time for every individual tree (Fig 1).

**Figure 1.**
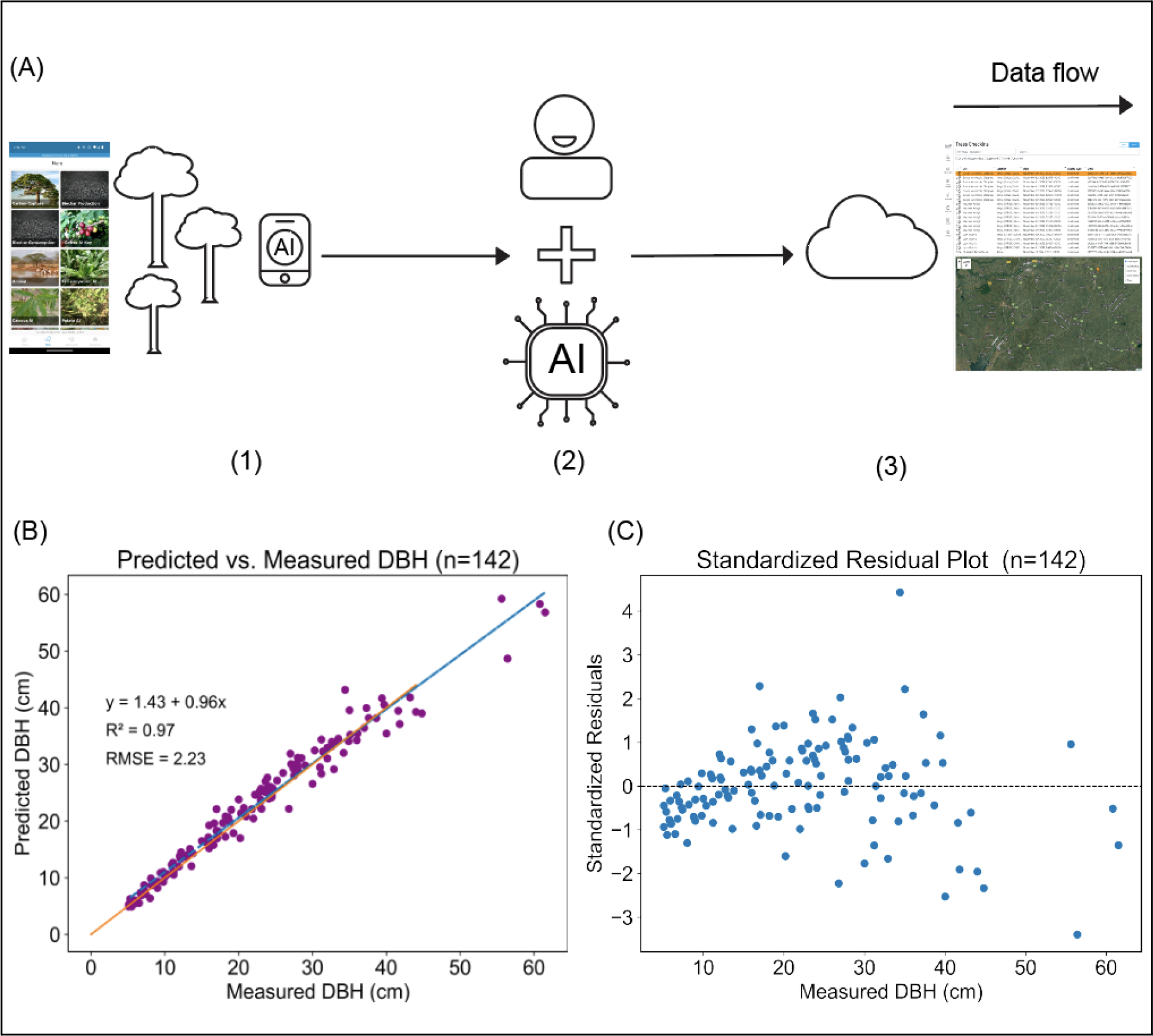
MRV System and the accuracy of the diameter estimation method that powers the MRV system. (A) The MRV System. (1) Farmers will use a mobile application (i.e., PlantVillage Nuru) to measure and report tree survival and growth over time. Farmers will measure trees by just taking a picture of the tree with a custom tree tag attached to the tree. (2) A human verifier with support from AI systems in the cloud will check and verify all tree info on a Web portal (i.e., PlantVillage Ag Observatory). (3) All verified agroforestry tree data can be publicly accessible on the Web portal. (B) Regression plot for evaluating the diameter measurement method developed in this study. The predicted and measured diameter values correlate with an R2 and RMSE of 0.97 and 2.23cm. (C) Standardized residual plot for the correlation analysis.

## 2 Materials and Methods

### 2.1 Data Collection

The open-source Open Data Kit mobile application was used to collect all the tree image datasets in this study (C Hartung et al., 2010). Data were collected from trees of various diameter values from 5 to 60 cm. For each tree, a measurement of its diameter and a picture of the tree with a custom tree tag were taken. A measuring tape was used for the diameter measurements of the trees. All diameter measurements were made at 1.37m above the ground. The tree tags were made from aluminum sheet metal with a QR code imprinted for the unique identification of each tree. The tree tags were made with aluminum sheets for durability purposes. The purpose of the unique tree tags is to link tree measurements and reports over time in the MRV system. A total of 192 trees were measured to calibrate (n=52) and evaluate (n=142) the diameter estimation method (Fig 2).

**Figure 2.**
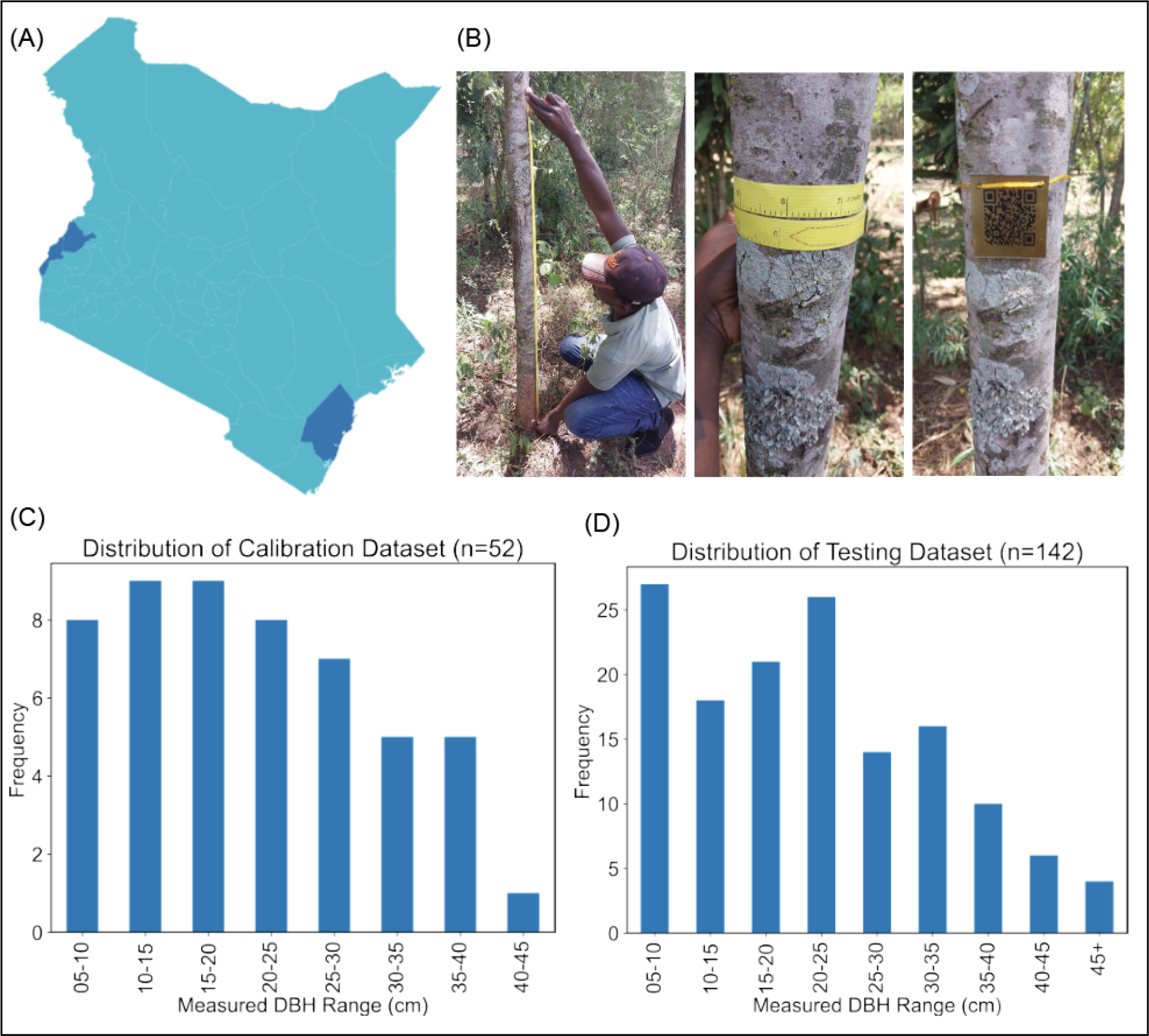
Data Collection. (A) Data was collected from Kenya’s Busia, Bungoma, and Kilifi counties. (B) Data was collected with the ODK mobile application. Images and a measurement of the tree’s diameter at 1.37m (diameter at breast height) were collected for each tree. All measurement values of trees were verified with a picture of the trees being measured. (C) Distribution of the dataset used to calibrate the diameter measurement method. (D) Distribution of dataset used to evaluate the diameter measurement method in this study.

An additional 494 images of trees with the custom tree tag were used to fine-tune the custom semantic segmentation model. The dataset was divided into 85% and 15% for training and evaluation datasets. All the 494 images were annotated with the PixelAnnotationTool (Bréhéret, 2017). PixelAnnotationTool enables annotators to assign a ground truth class label to every pixel in an image. The labels used for the dataset were background, tree trunk, and tag.

### 2.2 Segmentation Model Training and Evaluation

The semantic segmentation model was fine-tuned from the open-source DeepLab model from Google TensorFlow Research (LC Chen et al., 2018). The Deeplab model uses encoder-decoder structures in a deep neural network to perform image segmentation tasks. The former structures can encode multi-scale contextual information by probing the input image data with filters or probing operations at multiple rates and multiple effective fields of view, while the latter networks can capture sharper object boundaries by gradually recovering spatial information (LC Chen et al., 2018). The Deeplab model has achieved 89% and 82.1% test performance on the PASCAL VOC 2012 and Cityscapes datasets (LC Chen et al., 2018).

Transfer learning was applied to fine-tune a pre-trained semantic segmentation model on our dataset. The model’s architecture was Xception-65 and was pre-trained on the coco dataset (LC Chen et al., 2018). The segmentation model was trained for 300 epochs. The training batch size was 3. The initial learning rate was e^-4^ with a learning rate decay factor of 0.1 that began after step 6000. The atrous rates were [6,12,18], and the output stride was 16.

The performance of the segmentation model on the evaluation dataset after training was used for segmentation model evaluation. The mean and individual intersection over union (IOU) for the model labels reported after the final model evaluation was reported to quantify the performance of the segmentation model for classifying background, tree trunk, and tree tag pixels in an image. An IOU is between 0 and 1, where values close to 1 indicate that the model’s pixel classifications agree with the ground truth annotated pixels.

### 2.3 Diameter Estimation Algorithm

The diameter of trees was estimated after a series of tasks were operated on an image of the tree with a custom tree tag. The tree diameter estimation algorithm used in this experiment has six steps: (1) segmentation of input image, (2) check tree orientation, (3) cropping the image to increase the relative size of the tag, (4) segmentation of the cropped image, (5) calculate the tree-to-tag pixel width ratio, and finally, (6) estimate dbh (Fig 3).

**Figure 3.**
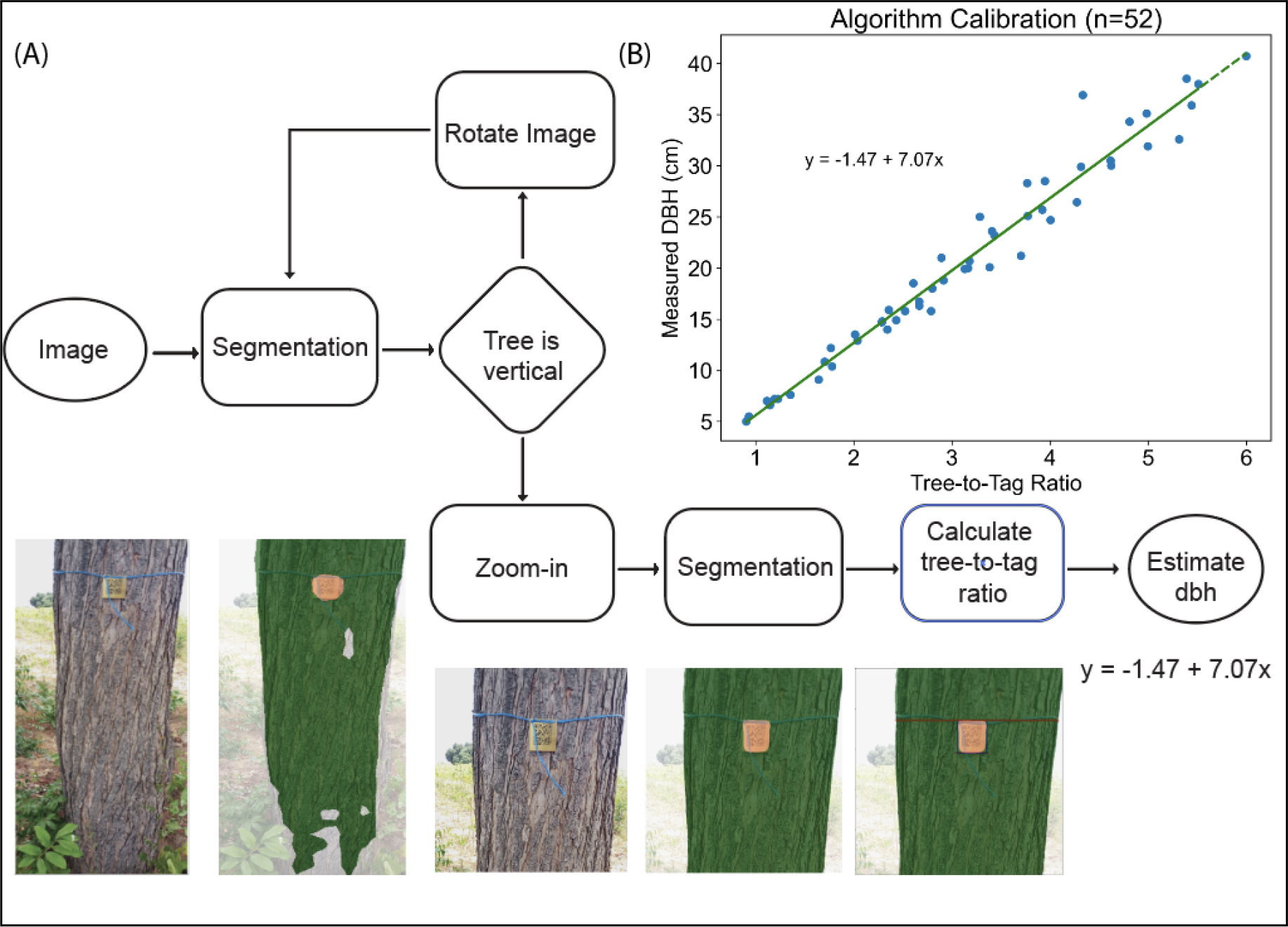
Diameter estimation algorithm. (A) The diameter estimation algorithm takes an image of a tree with the custom tree tag and predicts an accurate diameter. The algorithm generates two segmentation masks of the image. The algorithm uses the first segmentation mask to find the general location of the tag and zoom in close to the tag for a more precise image segmentation mask. The second mask is then used to estimate the tree-to-tag pixel width ratio, which is then used to estimate the tree’s diameter just above the tag. The tree’s diameter is calculated with the regression model trained to predict tree diameter based on the tree-to-tag pixel width ratio. (B) The regression model equation is y=-1.47 + 7.07x, where y is the tree’s diameter and x is the tree-to-tag pixel width ratio (n=52).

1. The fine-tuned segmentation model generates a segmentation mask from the input image. The model classifies each input image pixel into one of three categories: background, tag, and tree trunk. The segmentation mask is saved for the next step.
2. The algorithm expects the orientation of the tree in the images to be vertical. The algorithm checks and rotates images to ensure that the orientation of trees in all the images is vertical. The algorithm checks the trees’ orientation by checking if the detected tree pixels are oriented left to right or top to bottom.
3. The algorithm uses the segmentation mask from step (1) to find the general location of the tree tag in the image and zoom in. The algorithm finds the edges or boundaries of the tree tag and estimates the tree tag pixel width from the segmentation mask. The algorithm buffers pixels equal to the tree tag pixel width on the left and right sides of the detected tree edges or boundaries and at least two tree tag pixel widths above and below the detected tree tag in the image. The tree and tree tag edge detection and tree tag pixel width estimation are made using canny detection, contour detection, and minAreaRect image processing functions from the Python OpenCV library (OpenCV, 2015).
4. Similar to step (1), a semantic segmentation mask is generated for the zoomed image. The segmentation masks of the zoomed images were better for the edge detection of the tree and the tree tag than the segmentation masks on the original images. This is because of the inherent downsampling of the input image to a fixed shape before model inference. The zoomed images are less downsampled than the original large images, thus providing the model with high-quality images to generate inference. Accurate edge detection of the tree tag is critical for accurate diameter estimation. The generated segmentation mask is saved for the next step.
5. The algorithm calculates the tree pixel width and the tree tag pixel width to estimate a tree-to-tag pixel width ratio. The tree pixel width is estimated by calculating the average number of the tree trunk pixels on adjacent rows just above or below the detected tree tag. The number of pixel rows selected above or below the tree tag is equivalent to the tree tag pixel width. By default, the algorithm estimates the tree pixel width by averaging over the rows above the tag. However, in some cases, it will average the rows below the tag if the number of pixel rows above the tag is less than the tree tag pixel width. The tree tag pixel width is estimated, as explained in step 3. After estimating the tree and tree tag pixel widths, the algorithm calculates the tree-to-tag pixel width ratio and moves to the next step.
6. The algorithm estimates the diameter of a tree with a linear regression model trained to predict diameter based on the tree-to-tag pixel width ratio. The equation for the linear regression model is y = -1.47 + 7.07x, where y is the diameter, and x is the tree-to-tag ratio of the tree. The linear regression model was developed with ground truth diameter tape measurements and the tree-to-tag ratio of 52 annotated trees with custom tree tag images (Fig 3). This calibration enables the algorithm to overcome the photogrammetric error associated with image distortions from the phone camera lens (Chandrasekhar et al., 2011; Neale et al., 2011; Luhmann et al., 2016)

### 2.4 Diameter Estimation Algorithm Statistical Evaluation

The relationship between the predicted and measured diameter values was evaluated with correlation analysis. The performance of the regression line was evaluated with the coefficient of determination (R^2^), root means square error (RMSE), and a significance level of 0.05. The regression line of best fit and the standardized residual values were generated using the numpy and statsmodels libraries in Python (add version information). The RMSE was calculated with the equation below, where n is the sample size, predicted is the predicted diameter values, and measured is the measured diameter values. The R^2^ was calculated with the sklearn library in Python.

The performance of the measurement method for different diameter ranges was further investigated with bias (BIAS), relative bias (relBIAS), root means square error (RMSE), and relative root means square error (relRMSE) (Li et al., 2023). The diameter ranges were (1) 5.0 cm to 15.9 cm, (2) 16.0 cm to 25.9 cm, (3) 26.0 cm to 35.9 cm, and (4) 36 cm and above. BIAS calculates how well the predicted diameter values fit the measured diameter values - Eqn 1. The relBIAS is the absolute deviation of a measurement as a percentage of the mean - Eqn 2. The relBIAS intuitively shows the magnitude and direction of bias across the different diameter ranges. The RMSE is calculated by the Equation (3). The relRMSE was calculated with Equation (4). The relRMSE intuitively shows the different accuracy of the measurement method for the different diameter ranges.

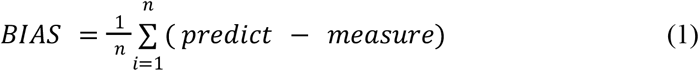

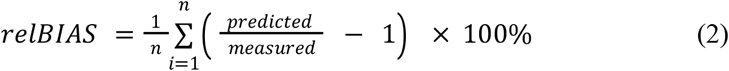

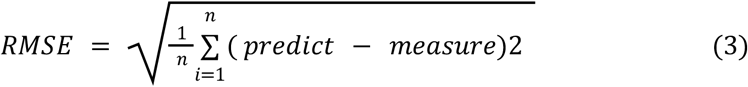

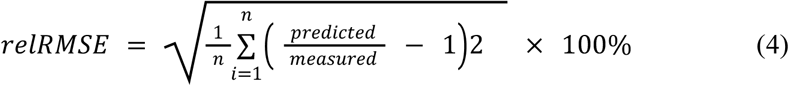

## 3 Results

### 3.1 Segmentation Model Evaluation Results

The segmentation model achieves an overall mean intersection over union of 0.937 over the evaluation dataset after model training (Table 1). The intersection over union of the background, tree trunk, and tree tag labels were 0.964, 0.930, and 0.918, respectively. The high IOU values reported here indicate that the segmentation model is accurate and reliable for background, tree trunk, and tree tag segmentation tasks.

**Table 1.** The evaluation of the segmentation model achieved an overall mean intersection over union (MIOU) of 0.937. The individual label IOU was 0.964, 0.930, and 0.918 for background, tree trunk, and tree tag labels. The segmentation model was trained and evaluated with 494 images (n=494).

| Label | IOU |
| --- | --- |
| Background | 0.964 |
| Tree Trunk | 0.930 |
| Tree Tag | 0.918 |
| MIOU | 0.937 |

### 3.2 Diameter Estimation Algorithm Evaluation Results

The regression analysis results indicate that the predicted diameter values compare with the measured diameter values with an R^2^ of 0.97 and RMSE of 2.23 (n=142) (Figure 1). The intercepts and slope for the regression model are both statistically significant at a significance level of 0.05 (Table 2). The slope and intercepts of the regression model are 0.958 and 1.434, respectively (Table 2).

**Table 2.** Parameter estimates and statistical fitting summary model for tree diameter estimation algorithm evaluation(n=142).

| Variable | Estimates | Standard Error | t-value | p > \|t\| |
| --- | --- | --- | --- | --- |
| Intercepts | 1.4344 | 0.369 | 3.885 | 0.000 |
| Measured diameter | 0.9579 | 0.015 | 65.150 | 0.000 |

The BIAS, relBIAS, RMSE, and relRMSE results (Table 3) indicate that the performance of the developed measurement method is variable across different diameter ranges. The BIAS and relBIAS reported for diameter measurements within 5cm to 35.9 cm are positive, whereas the BIAS and relBIAS for 36+ are negative. This indicates that the measurement method is biased to overestimate trees with the diameter of trees smaller than 36 cm and underestimate the diameter of trees equal to and above 36 cm. On the other hand, the RMSE increases as diameter ranges increase, whereas relRMSE decreases. This decrease in relRMSE as diameter ranges increase indicates that the larger the tree, the higher the accuracy of diameter estimation.

**Table 3.** The BIAS, relBIAS, RMSE, and relRMSE results are presented below. The results reported in this table indicate that the performance of the measurement method is variable across different diameter ranges.

| Range (cm) | Num | BIAS (cm) | relBIAS (%) | RMSE (cm) | relRMSE (%) |
| --- | --- | --- | --- | --- | --- |
| 5.0 - 15.9 | 49 | 0.465 | 3.952 | 1.012 | 10.232 |
| 16.0 - 25.9 | 44 | 1.145 | 5.609 | 2.029 | 10.294 |
| 26.0 - 35.9 | 31 | 0.989 | 3.334 | 2.864 | 9.355 |
| 36+ | 18 | -1.746 | -3.657 | 3.517 | 7.443 |
| 5.0 - 60 | 142 | 0.51 | 3.366 | 2.233 | 9.751 |

## 4 Discussion

Methods for diameter estimation with innovative technology have been investigated by many authors (Li et al., 2023; Koreň et al., 2017; Woo et al., 2021; Shao et al., 2022; Tran et al., 2023; Putra et al., 2021). Contact measurement methods with customized hand-held technologies provide some of the best accuracies with RMSE ranging from 0.33 to 0.52 cm (Li et al., 2023). Non-contact and non-reference measurement method with various sensors and post-processing software provides varying accuracy, with RMSE ranging from 0.77 to 54.38 cm (Koreň et al., 2017; Woo et al., 2021; Shao et al., 2022; Tran et al., 2023; Putra et al., 2021). These innovative contact and non-contact non-reference measurement tools can provide a reliable tool to automate tree diameter measurement and reporting, but their implementation can be costly due to their requirement for custom sensors. Thus making them inaccessible and impractical for developing an MRV for smallholder farmers.

Conversely, the non-contact reference-based measurement method can be more accessible to farmers as it only requires a smart mobile phone. However, the unreliable and inaccurate identification of tree trunks and reference objects with image processing techniques can make reference-based methods unreliable (Shao et al., 2022). Megalingam et al. (2022) used different image processing methods to identify tree trunks and reference objects for coconut tree diameter estimation and achieved an accuracy ranging from 70.04% to 83.63%. The robustness and accuracy of non-contact reference-based depends on the method used to identify and extract the boundaries of the tree trunk and the reference object. We have used a state-of-the-art deep-learning semantic segmentation model to overcome this challenge. This study’s fine-tuned semantic segmentation model achieved an MIOU of 0.937 on the evaluation dataset after training (Table 1). Evaluation of the measurement method on 142 trees achieved an R^2^, BIAS, and RMSE of 0.97, 0.51 cm, and 2.23 cm, respectively (Table 3). The measurement accuracy reported here is comparable with some of the most recent tree diameter measurement methods (Table 4). This is the first study to use semantic segmentation and image-processing methods to develop a non-contact reference-based tree diameter measurement and verification method. This measurement method can automate tree measurement, reporting, and verification for an MRV platform.

**Table 4.**
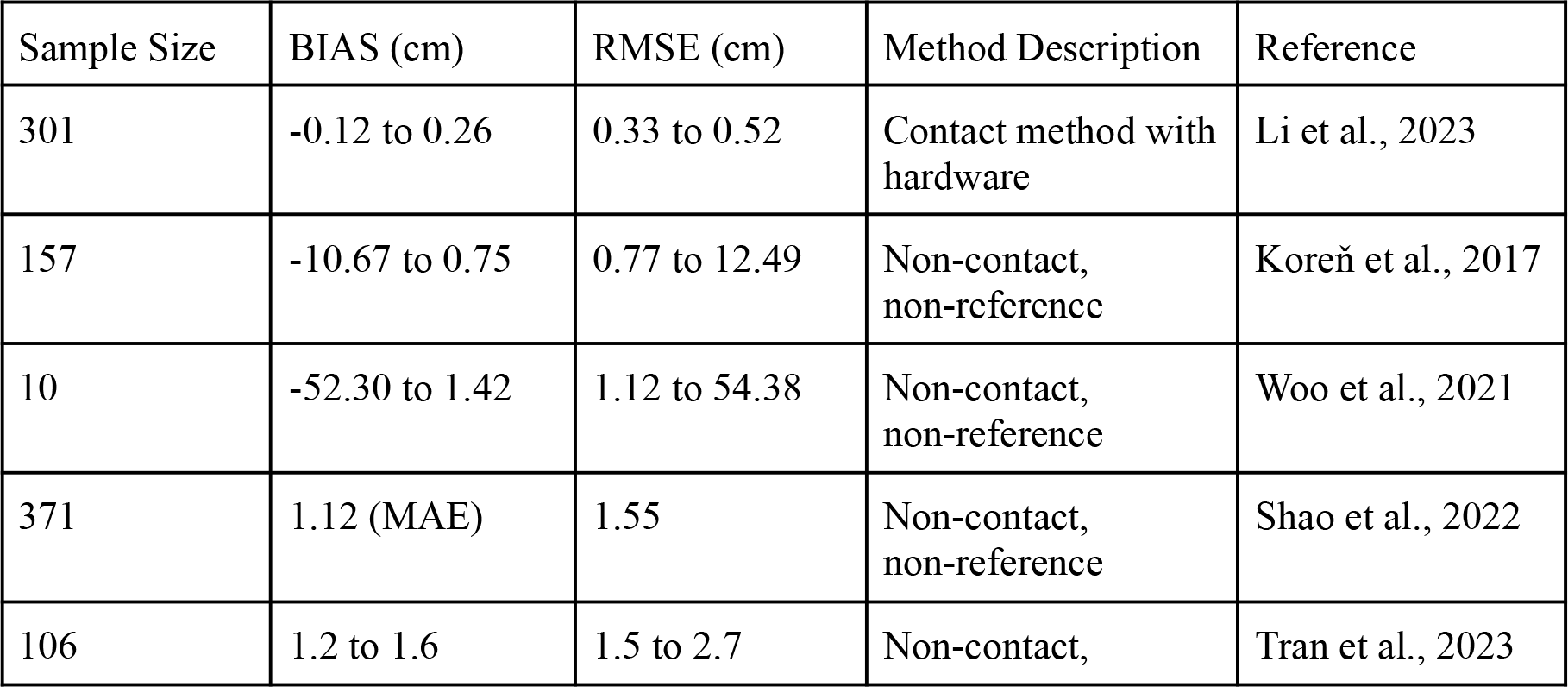
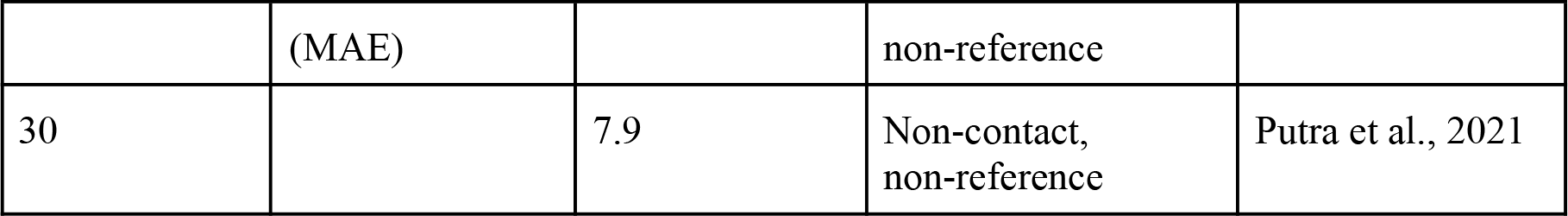
The BIAS and RMSE for other diameter estimation methods.

A detailed investigation of performance on different diameter ranges reveals that the method tends to overestimate and underestimate small and larger trees, respectively. One of the reasons for overestimating smaller trees is due to the algorithm’s sensitivity to the image’s semantic segmentation. Imprecise segmentation of the edges of the tree and the tree tag can result in overestimation, especially for smaller trees. The error associated with projecting a large 3D tree onto a 2D plane for the measure can result in underestimation due to photogrammetric phenomena like parallax and perspective distortion, especially when the image is taken close to the object (Jhan et al., 2017). Another limitation of the method here is the reference object itself. The upfront cost of making, distributing, and attaching the tree tags to the tree can be challenging. Another major limitation of this method is that the algorithm measures diameter just above the tree tag, so as the tree grows, the estimated diameter measurement will not be an accurate dbh value.

Future studies can improve this non-contact reference-based measurement method by improving the semantic segmentation model’s accuracy and the algorithm’s robustness. The latter will be more impactful. Improving the semantic segmentation model, especially for the boundaries between the tree and background, as well as the tree and the tree tag, can improve edge or boundary detection accuracy. The current algorithm does not consider the angle between the camera and the tree. However, this was not an issue for the dataset in this study as all images were taken with the phone camera relatively parallel to the tree and the tree tag, but this could be an issue during production with smallholder farmers. The current algorithm also doesn’t consider the distance between the tree and the phone camera. Machine learning methods for depth estimation (Mertan et al., 2022) can also improve diameter estimation, especially for larger trees, by accounting for the angle of view. The algorithm could be adjusted to account for the vertical position of the tree tag from the ground so that diameter measurement can be adjusted accordingly to measure the diameter at breast height.

The measurement, reporting, and verification (MRV) system described in this study can empower organizations or institutions to plant, grow, and monitor tree survival and growth with anyone, including smallholder farmers, with just a smartphone and an internet connection. Rosenstock et al. (2019) found that many non-Annex I or developing countries will use agroforestry for climate adaptation and mitigation. The study also found that the lack of reliable MRV for agroforestry trees in developing countries is one of the biggest challenges for scaling this climate mitigation and adaptation effort. Low-cost, accurate, non-contact approaches for measuring tree biomass are needed to scale climate-smart efforts with smallholder farmers (Ewing et al., 2021; Schilling et al., 2023). The MRV system described in this study can provide a low-cost, accurate, and reliable MRV for implementing tree planting efforts with anyone, including smallholder farmers.

## 8 Author Contributions

Edward Amoah and David Peter Hughes conceptualized the study. Edward Amoah performed the investigation in the study. David Peter Hughes, Peter McCloskey, Rimnoma Ouedrago, and John Chelal supervised the study. Chelsea Akuleut, Binti Ibrahan Mwambumba, Brain Kipchirchir Meli, Christabel Akinyi Oyudi, Edna Santa Kibwamga, Eunice Kwamboka Cleopha, Fredrick Odhiambo Ochola, Catherine Njeri Wangiru, Kelvin Morang’a, Lyon Wilson Mushira, Maureen Kalegi Maboke, Nancy Syonthi Titus, Serah Lanoi Oltimbao, and Sheilah Awour Odawa curated data. Edward Amoah trained the machine learning model in this study with supervision and guidance from Peter McCloskey. Edward Amoadh wrote the code for the algorithm. Edward Amoah performed the formal analysis of the study. Edward Amoah wrote the original draft of the study. David Peter Hughes, Peter McCloskey, Rommona Ouedrago, and Edward Amoah reviewed and edited the paper.

## 10 Acknowledgments

The work was supported by funds from the Elon Musk Foundation XPrize Carbon Removal Student Award. The Google TensorFlow Research DeepLab2 project was critical to developing the semantic segmentation model in this article. Thanks to the PlantVillage Lab for supporting this project’s data collection and implementation. Thanks to the numerous Penn State University undergraduates who annotated the tree images to develop this article. Thanks to the Dream Team Agro Consultancy members for helping with data collection.

## 12 Data Availability Statement

The data generated and analyzed for this study can be found on the git repository (https://github.com/eai6/pv_mrv/tree/ac1a5a5181094cc0fc4a3af7d111e1e30c38b180/statistical_analysis/output). The image dataset that was used to calibrate and evaluate the algorithm can be found on the ScholarShpere repository of the Pennsylvania State University (https://scholarsphere.psu.edu/resources/08a985a4-d878-4fa9-b2f2-6060100580a9). The diameter estimation tool calibrated for the custom tag used in this study is containerized and can be accessed publicly at this docker link (https://hub.docker.com/r/eai6/pv_mrv_api).

